# Synonymous lysine codon usage modification in a mobile antibiotic resistance gene similarly alters protein production in bacterial species with divergent lysine codon usage biases because it removes a duplicate AAA lysine codon

**DOI:** 10.1101/294173

**Authors:** Mohammed Alorabi, Aisha M. AlAmri, Yuiko Takebayashi, Kate J. Heesom, Matthew B. Avison

**Author notes:** Correspondence: Matthew B. Avison, School of Cellular & Molecular Medicine, Biomedical Sciences Building, University Walk, Bristol. BS81TD. UK.

## Abstract

The mobile antibiotic resistance gene *bla*_IMP-1_ is clinically important and has a synonymous AAA:AAG lysine codon usage bias of 73:27. This bias is like that seen in experimentally determined highly expressed genes in *Escherichia coli* and *Acinetobacter baumanii*, but quite different from that seen in *Pseudomonas aeruginosa* (26:74 AAA:AAG). Here we show that, paradoxically, shifting the AAA:AAG lysine codon bias to 8:92 in *bla*_IMP-1_ expressed from a natural promoter results in significantly more IMP-1 production in all three species. Sequential site directed mutagenesis revealed that increased IMP-1 production occurs following removal of an AAA,AAA double lysine codon and that otherwise, lysine codon usage had no observable impact on IMP-1 production. We conclude that ribosomal slippage at this poly-adenosine region reduces efficient translation of IMP-1 and that punctuating the region with guanine reduces ribosomal slippage and increases IMP-1 production.

## Introduction

Synonymous codon usage bias (SCUB) is a term describing the common finding that organisms favour the use of certain triplet codons in DNA to encode certain amino acids. Since SCUB varies between organisms, and between different genes in a single organism, the implication is that optimal SCUB varies between different organisms, and that certain genes are selected to be closer to the optimal SCUB than others (1). One dominant hypothesis is that highly expressed genes have “optimised” SCUB and that this is selected because optimal codons are translated more quickly and/or more efficiently than sub-optimal ones. This is particularly important when the demand for a protein is high (1). Indeed, it is well known that SCUB optimization – adapting the SCUB of a recombinant gene to match that of highly expressed genes – increases recombinant protein production in a heterologous host (2). The success of this methodological approach has been used to advance the translationally-selective hypothesis to explain SCUB. However, most codon usage optimization procedures involve the over-expression of recombinant genes using hyper-strong, inducible promoters and high copy number vectors, with a desire to make a single protein represent a high percentage of total protein in the cell (3). This is not likely to reflect the situation encountered by a gene in a natural setting with a natural promoter on the chromosome or a low copy number plasmid.

SCUB is particularly relevant in the context of horizontal gene transfer. Whilst sub-optimal SCUB is not always seen in horizontally acquired genes, depending on their origins, for those that are sub-optimal, selective pressure is expected to be applied over time to optimise SCUB; a process referred to as “codon usage amelioration” (4). Antibiotic resistance in bacteria is one of the most pressing threats to human health and horizontal gene transfer is one of the most important means for a bacterium to acquire antibiotic resistance (5). It is evident that many mobile genetic elements are of low guanine plus cytosine (GC) content and carry a related SCUB biased towards low GC codons (6). So, in this context, the impact of SCUB on gene expression is not only academic, but it is also of significant practical interest. The aim of the work reported in this paper was to test the hypothesis that SCUB change can affect the absolute amount of active protein produced in a predictable manner using a natural plasmid system and an intermediate strength natural promoter using a clinically important, mobile antibiotic resistance gene.

## Results and Discussion

The *bla*_IMP-1_ metallo-β-lactamase-encoding gene cassette, which confers resistance to the carbapenems, a class of “last resort” antibiotics, has a GC content of 39% (7). However, it has been found widely among Gram-negative bacteria with varying genomic GC contents (8). For example, it is commonly found in *Escherichia coli* and other Enterobacteriaceae which have genomic GCs of ≈50%. Accordingly, if SCUB affects translational efficiency and/or rate, one would expect the amount if IMP-1 enzyme should be seen to increase in Enterobacteriaceae if the SCUB of *bla*_IMP-1_ is ameliorated to match the optimal SCUB of this family, as represented by *E. coli*. To test this, specific imipenemase activity was measured in extracts of *E. coli* MG1655 transformants carrying pHIMP or pH*Ec*IMP, being the cloned wild-type or *E. coli* SCUB optimised *bla*_IMP-1_ gene cassette, respectively, each under the control of an identical, natural, intermediate-strength integron promoter, and each ligated into a broad host-range, low copy number vector derived from a natural antibiotic resistance plasmid: RK2 (9). We can confirm that pHIMP is a truly natural expression system because imipenemase activity in an MG1655 transformant carrying pNIMP (the natural *bla*_IMP-1_ encoding plasmid) and MG1655(pHIMP) were the same (**Figure 1**). If the hypothesis being tested is correct, it was expected that optimisation of *bla*_IMP-1_ to a SCUB closer to optimal in *E. coli* genes would increase IMP-1 production relative to wild-type. However, this was not observed. The pH*Ec*IMP variant synthesised to match the “optimal” SCUB of *E. coli* presumed highly expressed genes (ribosomal protein and translation elongation factor genes) based on the OPTIMIZER algorithm (10) was expressed at lower levels than than the wild-type gene (p<0.0001) (**Figure 1**).

**Figure 1:**
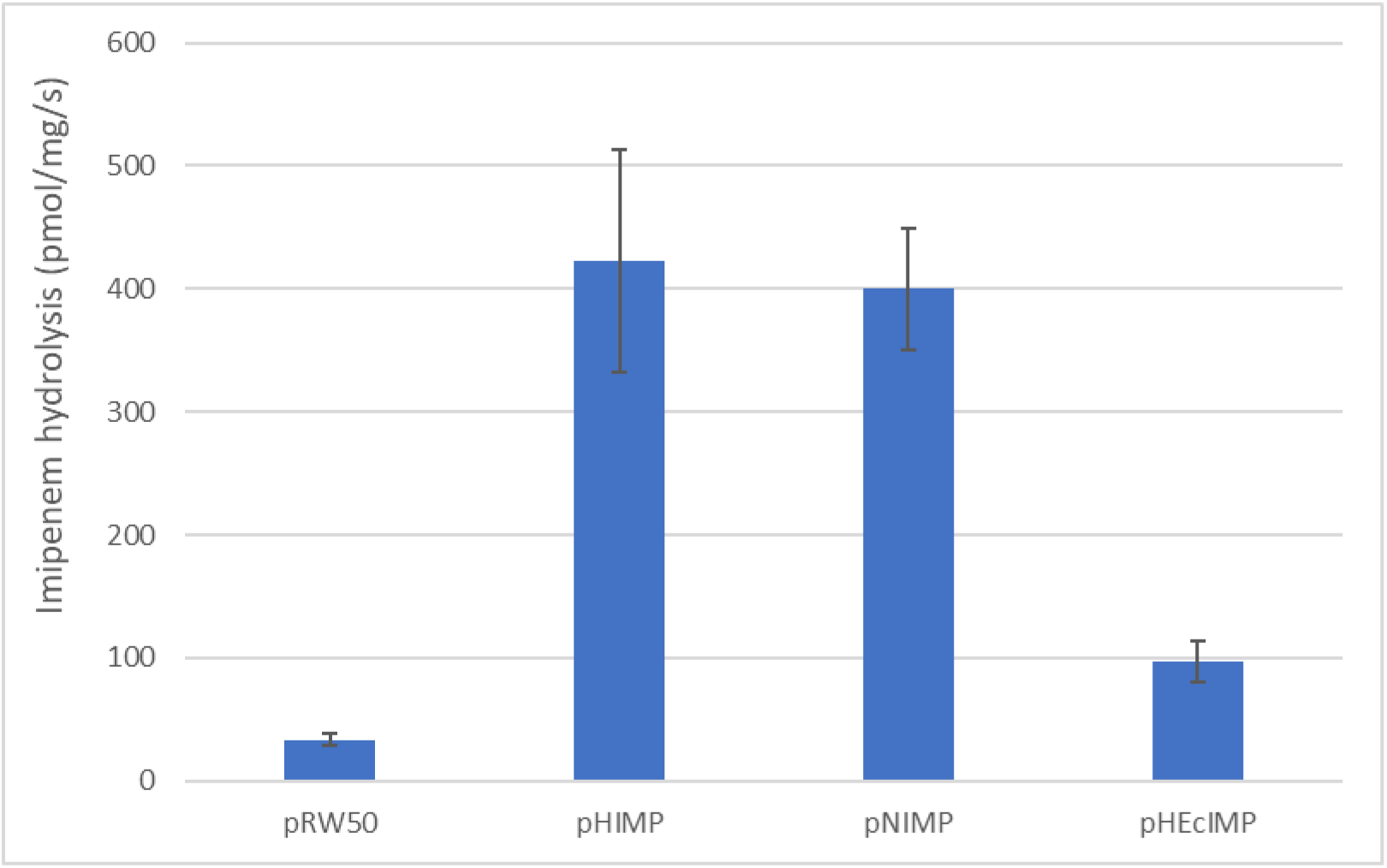
Impact of codon usage “optimisation” on IMP-1 enzyme activity in *E. coli*. IMP-1 enzyme activity was measured using imipenem as substrate in whole cell extracts of *E. coli* MG1655 carrying empty vector pRW50, or this vector with the wild-type (pHIMP) or codon-optimised (pHEcIMP) *bla*_IMP-1_ gene, each expressed from a hybrid strength integron promoter; or carrying a natural IMP-1 encoding plasmid originally from *P. aeruginosa*, which is the source of the cloned *bla*_IMP-1_ and upstream/promoter sequence (pNIMP). Data are means +/-Standard Deviation, *n*=4.

It was considered possible that the effect seen here is due to stability changes at the 5’ end of the codon-optimised *bla*_IMP-1_ mRNA. Strong regions of secondary structure at the 5’ ends of mRNA molecules are likely to cause ribosomal occlusion leading to the exposure of mRNA to nuclease digestion (11). Furthermore, it has previously been shown that synonymous mutations that increase mRNA secondary structure (high folding energy) at the beginning of the transcript can reduce protein production by inhibiting the initiation and initial phase of translation elongation (12-14). It was confirmed that codon optimization increased the energy required to unfold the mRNA. The Gibbs free energy value of the whole mRNA molecule was −183 (wild type *bla*_IMP-1_) changing to −229 for the *E. coli* codon optimised variant. Just looking at the 5’ third of the mRNA, which is thought to be particularly important, the folding energy calculated showed the same effect: moving from −50 for wild-type *bla*_IMP-1_ to −67 for the *E. coli* variant.

**Figure 1** shows evidence, therefore, that codon optimization can have negative effects on gene expression in a natural expression system, but it was considered of interest to see how much of a change in IMP-1 production would occur upon site-directed mutation of individual codons. Twenty six of 246 (10.6%) of IMP-1’s amino acids are lysine. The lysine codon AAA is in the majority in *bla*_IMP-1_ (19/26; 73%) and AAG accounts for the rest. Rather than relying on theoretical lists of “highly expressed genes” to define the “optimal” SCUB for lysine codons in *bla*_IMP-1_, we measured protein abundance using LC-MS/MS proteomics. In so doing we defined the 20 most highly abundant proteins in three test species during growth under the conditions we would also use to test IMP-1 production (**Tables 1-3**). Analysis of lysine SCUB in the genes encoding these 20 proteins from each species revealed an AAA percentage of 84% for *Acinetobacter baumannii*, 80% for *E. coli* and 24% for *Pseudomonas aeruginosa* (**Tables 4-6**). We sub-cloned the wild-type *bla*_IMP-1_ gene from pHIMP into the pSU18 cloning vector and used site directed mutagenesis to dramatically reduced the AAA lysine codon usage of *bla*_IMP-1_ in the resultant pSUHIMP-WT plasmid to 2/26 or 8% AAA in plasmid pSUHIMP-KV. Plasmids were used to transform an *E. coli* clinical isolate to chloramphenicol resistance. The wild-type and lysine codon-variant *bla*_IMP-1_ genes were also sub-cloned into the broad host-range vector pUBYT, generating pUBYTHIMP-WT and pUBYTHIMP-KV, which were used to transform *P. aeruginosa* PA01 or *A. baumannii* CIP 70-10 to kanamycin resistance. We then used proteomics to measure the abundance of IMP-1 in these transformants, which was normalised using the abundance of vector-encoded Cat (in *E. coli* pSUHIMP-WT and -KV transformants) or AphA (in *A. baumannii* and *P. aeruginosa* pUBYTHIMP-WT and -KV transformants) to take into consideration plasmid copy number and protein loading. Our expectation given the hypothesis that SCUB is selected based on translation rate or efficiency was that as AAA usage was reduced to 2/26 from a wild-type position of 19/26, which is close to optimal in *A. baumannii* (22/26) and *E. coli* (21/26) there would be a reduction in IMP-1 production. The case in *P. aeruginosa* was not so clear, given that the optimal AAA usage in this species is 6/26. Here, the variant is closer to optimal than the wild-type gene, so we might expect an increase in IMP-1 production. We did see this: a 1.5-fold increased normalised IMP-1 production in PA01(pUBYTHIMP-KV) compared with PA01(pUBYTHIMP-WT), *p*=0.04 for an unpaired t-test, *n*=3. However, the variant also supported higher IMP-1 protein production in *E. coli* (2.2-fold, *p*=0.005 *n*=3) and *A. baumannii* (3.2-fold, *p*=0.002, *n*=3) (**Figure 2**).

**Table 1.** The 20 most highly abundant proteins in *E. coli* during growth in Nutrient Broth

| Accession | Description | Gene Name | Mean Abundance (n=3) |
| --- | --- | --- | --- |
| P0CE48 | Elongation factor Tu | b3980 | 6.184E+08 |
| P0A9B2 | Glyceraldehyde-3-phosphate dehydrogenase A | b1779 | 3.399E+08 |
| P69776 | Major outer membrane lipoprotein Lpp | b1677 | 2.654E+08 |
| P0A7J3 | 50S ribosomal protein L10 | b3985 | 2.539E+08 |
| P06996 | Outer membrane protein C | b2215 | 2.425E+08 |
| P02359 | 30S ribosomal protein S7 | b3341 | 2.357E+08 |
| P0A7R1 | 50S ribosomal protein L9 | b4203 | 2.253E+08 |
| P0A7L0 | 50S ribosomal protein L1 | b3984 | 2.219E+08 |
| P60438 | 50S ribosomal protein L3 | b3320 | 2.022E+08 |
| P0AG55 | 50S ribosomal protein L6 | b3305 | 1.825E+08 |
| P0A910 | Outer membrane protein A | b0957 | 1.780E+08 |
| P0A7W1 | 30S ribosomal protein S5 | b3303 | 1.752E+08 |
| P62399 | 50S ribosomal protein L5 | b3308 | 1.743E+08 |
| P0A6M8 | Elongation factor G | b3340 | 1.644E+08 |
| P0A7V8 | 30S ribosomal protein S4 | b3296 | 1.642E+08 |
| P0A7R5 | 30S ribosomal protein S10 | b3321 | 1.622E+08 |
| P0AG67 | 30S ribosomal protein S1 | b0911 | 1.541E+08 |
| P0A7X3 | 30S ribosomal protein S9 | b3230 | 1.486E+08 |
| P0A7J7 | 50S ribosomal protein L11 | b3983 | 1.434E+08 |
| P0A7K2 | 50S ribosomal protein L7/L12 | b3986 | 1.422E+08 |

**Table 2:**
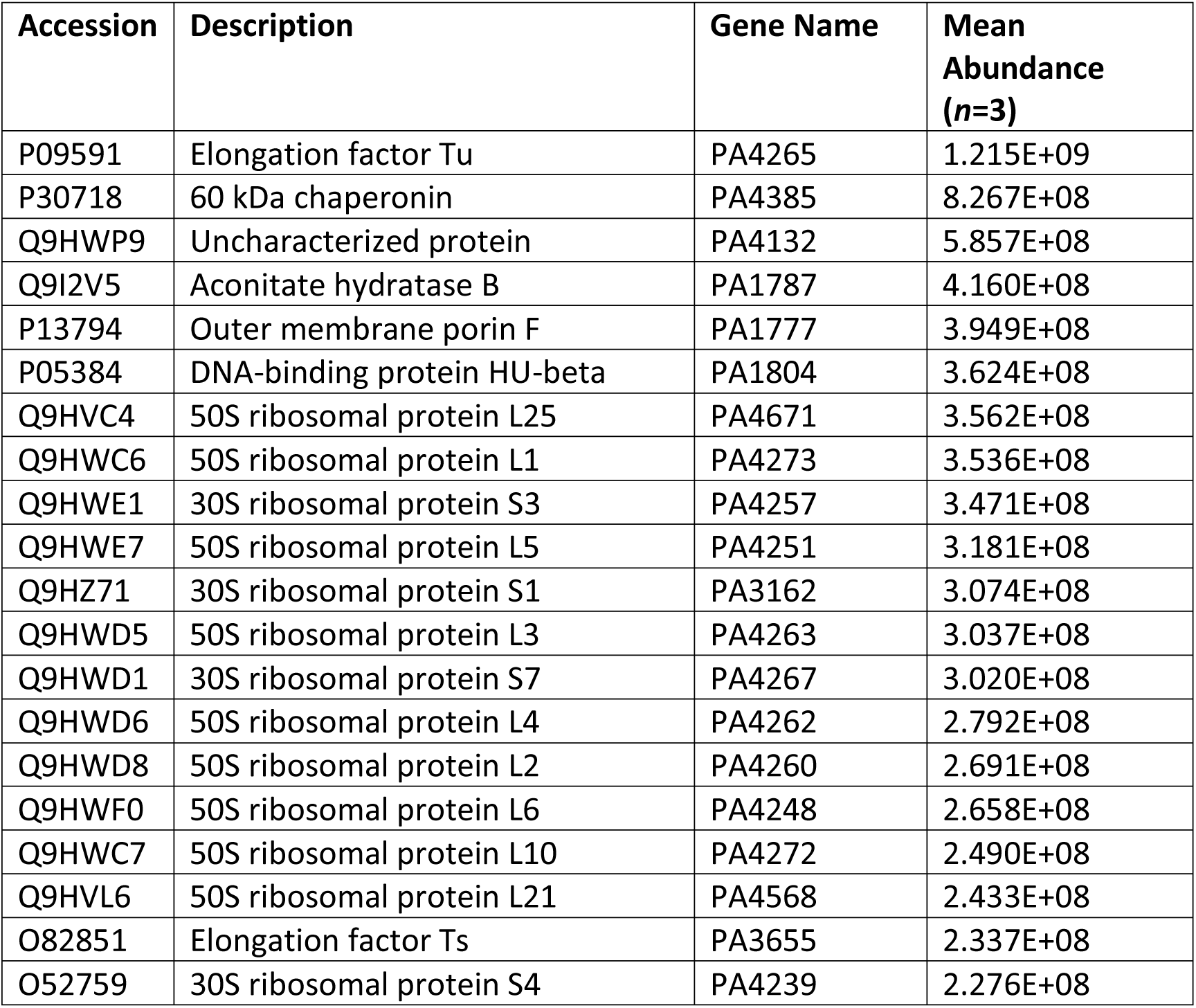
The 20 most highly abundant proteins in *P. aeruginosa* during growth in Nutrient Broth

**Table 3:**
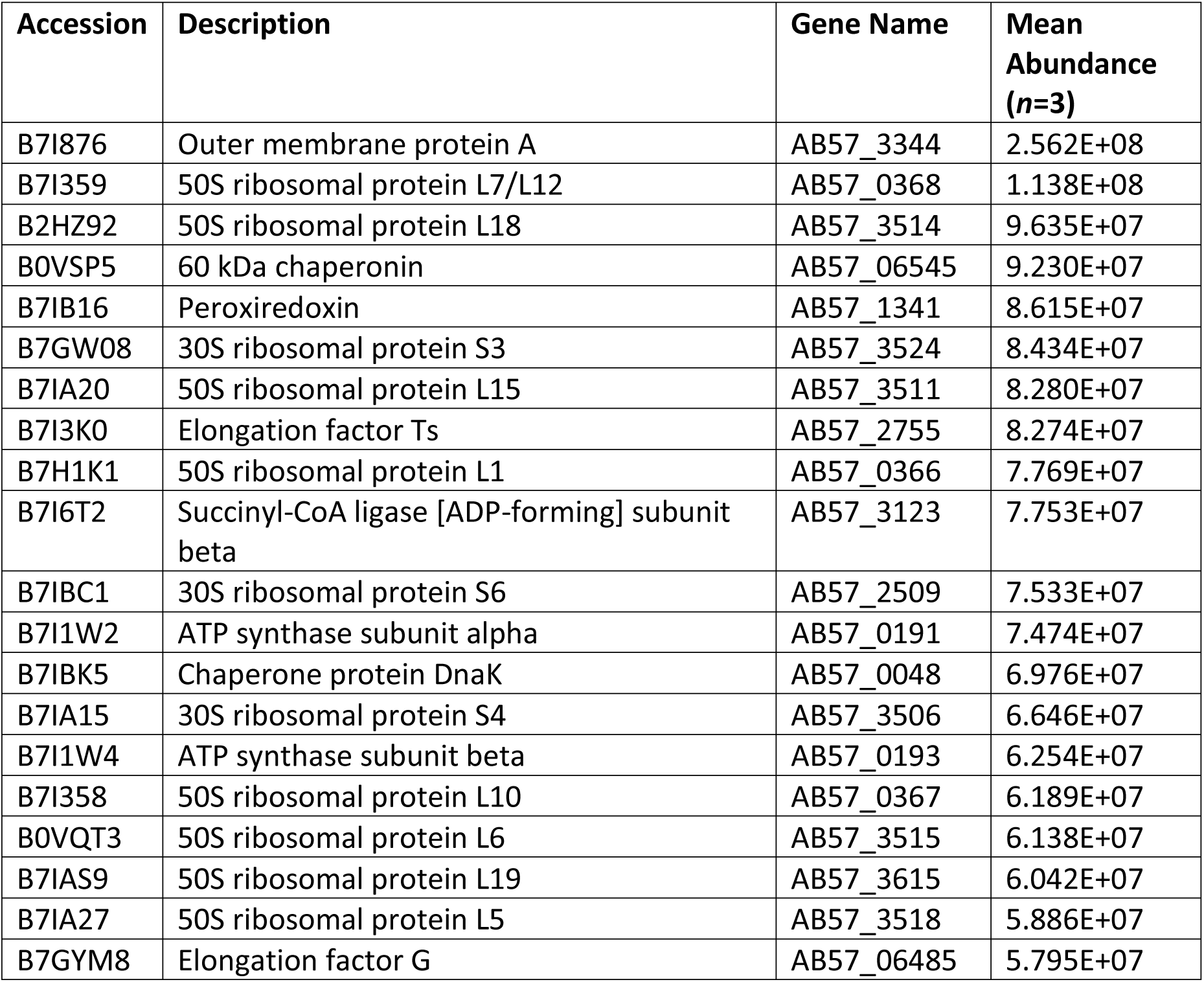
The 20 most highly abundant proteins in *A. baumannii* during growth in Nutrient Broth

**Table 4:**
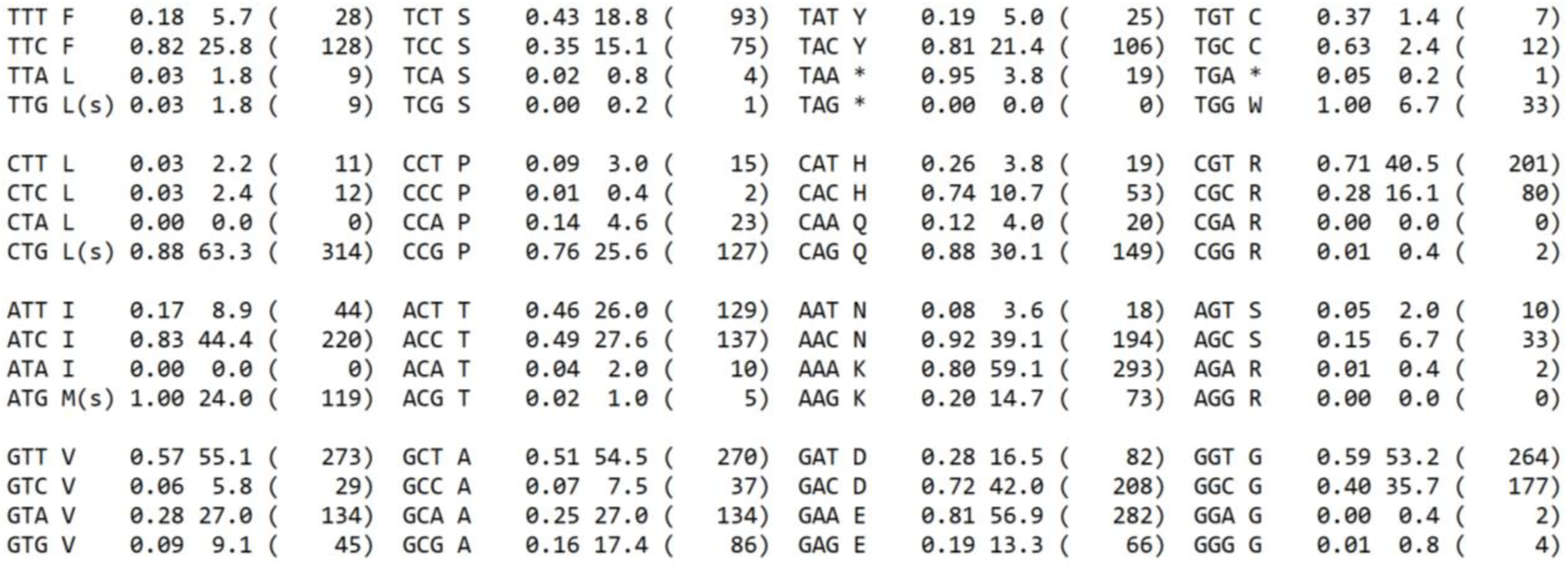
Codon Usage Table for 20 Most Highly Expressed Genes in *E. coli*

|  |  |  |  |  |  |  |  |  |  |  |  |  |  |  |  |  |  |  |  |
| --- | --- | --- | --- | --- | --- | --- | --- | --- | --- | --- | --- | --- | --- | --- | --- | --- | --- | --- | --- |
| TTT F | 0.18 | 5.7 | ( | 28) | TCT S | 0.43 | 18.8 | ( | 93) | TAT Y | 0.19 | 5.0 | ( | 25) | TGT C | 0.37 | 1.4 | ( | 7) |
| TTC F | 0.82 | 25.8 | ( | 128) | TCC S | 0.35 | 15.1 | ( | 75) | TAC Y | 0.81 | 21.4 | ( | 106) | TGC C | 0.63 | 2.4 | ( | 12) |
| TTA L | 0.03 | 1.8 | ( | 9) | TCA S | 0.02 | 0.8 | ( | 4) | TAA * | 0.95 | 3.8 | ( | 19) | TGA * | 0.05 | 0.2 | ( | 1) |
| TTG L(s) | 0.03 | 1.8 | ( | 9) | TCG S | 0.00 | 0.2 | ( | 1) | TAG * | 0.00 | 0.0 | ( | 0) | TGG W | 1.00 | 6.7 | ( | 33) |
| CTT L | 0.03 | 2.2 | ( | 11) | CCT P | 0.09 | 3.0 | ( | 15) | CAT H | 0.26 | 3.8 | ( | 19) | CGT R | 0.71 | 40.5 | ( | 201) |
| CTC L | 0.03 | 2.4 | ( | 12) | CCC P | 0.01 | 0.4 | ( | 2) | CAC H | 0.74 | 10.7 | ( | 53) | CGC R | 0.28 | 16.1 | ( | 80) |
| CTA L | 0.00 | 0.0 | ( | 0) | CCA P | 0.14 | 4.6 | ( | 23) | CAA Q | 0.12 | 4.0 | ( | 20) | CGA R | 0.00 | 0.0 | ( | 0) |
| CTG L(s) | 0.88 | 63.3 | ( | 314) | CCG P | 0.76 | 25.6 | ( | 127) | CAG Q | 0.88 | 30.1 | ( | 149) | CGG R | 0.01 | 0.4 | ( | 2) |
| ATT I | 0.17 | 8.9 | ( | 44) | ACT T | 0.46 | 26.0 | ( | 129) | AAT N | 0.08 | 3.6 | ( | 18) | AGT S | 0.05 | 2.0 | ( | 10) |
| ATC I | 0.83 | 44.4 | ( | 220) | ACC T | 0.49 | 27.6 | ( | 137) | AAC N | 0.92 | 39.1 | ( | 194) | AGC S | 0.15 | 6.7 | ( | 33) |
| ATA I | 0.00 | 0.0 | ( | 0) | ACA T | 0.04 | 2.0 | ( | 10) | AAA K | 0.80 | 59.1 | ( | 293) | AGA R | 0.01 | 0.4 | ( | 2) |
| ATG M(s) | 1.00 | 24.0 | ( | 119) | ACG T | 0.02 | 1.0 | ( | 5) | AAG K | 0.20 | 14.7 | ( | 73) | AGG R | 0.00 | 0.0 | ( | 0) |
| GTT V | 0.57 | 55.1 | ( | 273) | GCT A | 0.51 | 54.5 | ( | 270) | GAT D | 0.28 | 16.5 | ( | 82) | GGT G | 0.59 | 53.2 | ( | 264) |
| GTC V | 0.06 | 5.8 | ( | 29) | GCC A | 0.07 | 7.5 | ( | 37) | GAC D | 0.72 | 42.0 | ( | 208) | GGC G | 0.40 | 35.7 | ( | 177) |
| GTA V | 0.28 | 27.0 | ( | 134) | GCA A | 0.25 | 27.0 | ( | 134) | GAA E | 0.81 | 56.9 | ( | 282) | GGA G | 0.00 | 0.4 | ( | 2) |
| GTG V | 0.09 | 9.1 | ( | 45) | GCG A | 0.16 | 17.4 | ( | 86) | GAG E | 0.19 | 13.3 | ( | 66) | GGG G | 0.01 | 0.8 | ( | 4) |

**Table 5:** Codon Usage Table for 20 Most Highly Expressed Genes in *P. aeruginosa*

|  |  |  |  |  |  |  |  |  |  |  |  |  |  |  |  |  |  |  |  |
| --- | --- | --- | --- | --- | --- | --- | --- | --- | --- | --- | --- | --- | --- | --- | --- | --- | --- | --- | --- |
| TTT F | 0.08 | 2.1 | ( | 11) | TCT S | 0.04 | 1.9 | ( | 10) | TAT Y | 0.07 | 1.4 | ( | 7) | TGT C | 0.03 | 0.2 | ( | 1) |
| TTC F | 0.92 | 25.6 | ( | 132) | TCC S | 0.48 | 23.3 | ( | 120) | TAC Y | 0.93 | 18.8 | ( | 97) | TGC C | 0.97 | 5.2 | ( | 27) |
| TTA L | 0.00 | 0.2 | ( | 1) | TCA S | 0.00 | 0.0 | ( | 0) | TAA * | 0.75 | 2.9 | ( | 15) | TGA * | 0.20 | 0.8 | ( | 4) |
| TTG L(s) | 0.01 | 1.2 | ( | 6) | TCG S | 0.16 | 7.8 | ( | 40) | TAG * | 0.05 | 0.2 | ( | 1) | TGG W | 1.00 | 5.0 | ( | 26) |
| CTT L | 0.01 | 1.0 | ( | 5) | CCT P | 0.08 | 2.7 | ( | 14) | CAT H | 0.24 | 4.3 | ( | 22) | CGT R | 0.44 | 29.3 | ( | 151) |
| CTC L | 0.12 | 9.7 | ( | 50) | CCC P | 0.17 | 6.2 | ( | 32) | CAC H | 0.76 | 13.4 | ( | 69) | CGC R | 0.50 | 33.0 | ( | 170) |
| CTA L | 0.00 | 0.4 | ( | 2) | CCA P | 0.01 | 0.2 | ( | 1) | CAA Q | 0.15 | 5.2 | ( | 27) | CGA R | 0.01 | 0.6 | ( | 3) |
| CTG L(s) | 0.84 | 66.5 | ( | 343) | CCG P | 0.75 | 27.0 | ( | 139) | CAG Q | 0.85 | 29.3 | ( | 151) | CGG R | 0.04 | 2.7 | ( | 14) |
| ATT I | 0.09 | 4.8 | ( | 25) | ACT T | 0.12 | 5.2 | ( | 27) | AAT N | 0.08 | 2.5 | ( | 13) | AGT S | 0.02 | 1.2 | ( | 6) |
| ATC I | 0.91 | 48.3 | ( | 249) | ACC T | 0.85 | 37.1 | ( | 191) | AAC N | 0.92 | 30.5 | ( | 157) | AGC S | 0.30 | 14.4 | ( | 74) |
| ATA I | 0.00 | 0.0 | ( | 0) | ACA T | 0.01 | 0.4 | ( | 2) | AAA K | 0.24 | 17.1 | ( | 88) | AGA R | 0.00 | 0.2 | ( | 1) |
| ATG M(s) | 1.00 | 23.3 | ( | 120) | ACG T | 0.02 | 0.8 | ( | 4) | AAG K | 0.76 | 55.5 | ( | 286) | AGG R | 0.00 | 0.2 | ( | 1) |
| GTT V | 0.20 | 20.0 | ( | 103) | GCT A | 0.27 | 29.5 | ( | 152) | GAT D | 0.22 | 11.6 | ( | 60) | GGT G | 0.35 | 32.2 | ( | 166) |
| GTC V | 0.42 | 42.1 | ( | 217) | GCC A | 0.51 | 55.7 | ( | 287) | GAC D | 0.78 | 41.9 | ( | 216) | GGC G | 0.62 | 56.7 | ( | 292) |
| GTA V | 0.12 | 12.0 | ( | 62) | GCA A | 0.08 | 8.3 | ( | 43) | GAA E | 0.57 | 43.1 | ( | 222) | GGA G | 0.01 | 1.0 | ( | 5) |
| GTG V | 0.27 | 26.8 | ( | 138) | GCG A | 0.14 | 15.1 | ( | 78) | GAG E | 0.43 | 32.6 | ( | 168) | GGG G | 0.02 | 1.9 | ( | 10) |

**Table 6:**
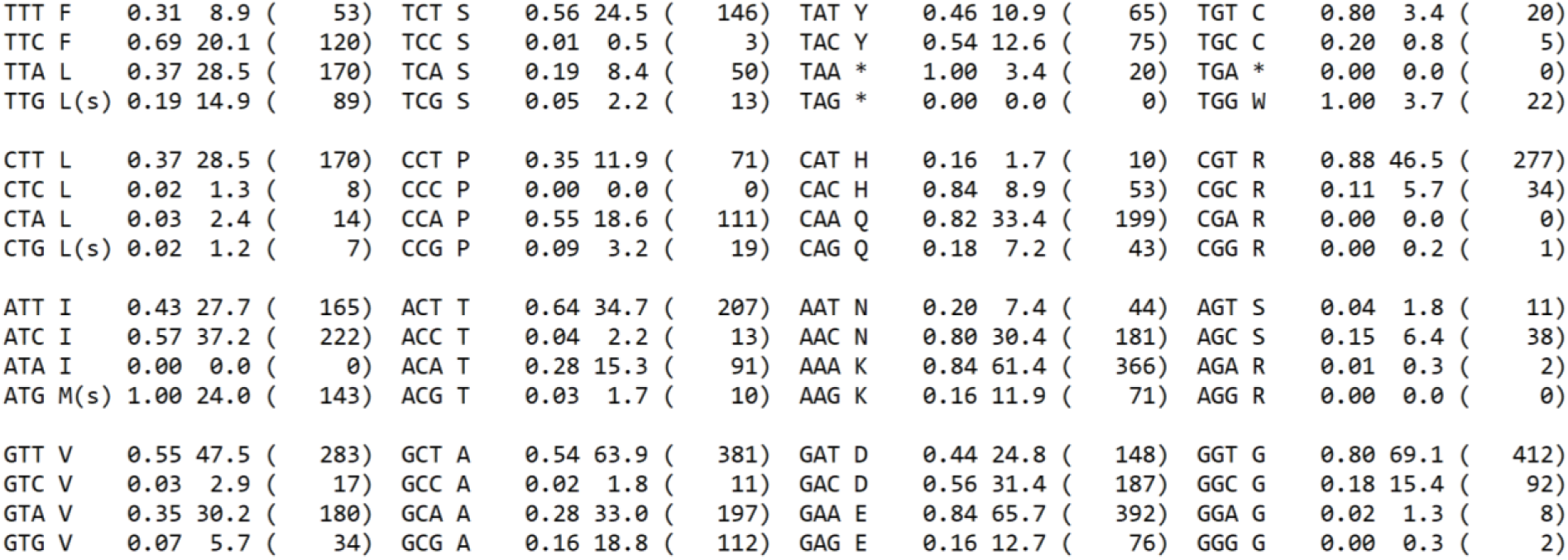
Codon Usage Table for 20 Most Highly Expressed Genes in *A. baumannii*

|  |  |  |  |  |  |  |  |  |  |  |  |  |  |  |  |
| --- | --- | --- | --- | --- | --- | --- | --- | --- | --- | --- | --- | --- | --- | --- | --- |
| TTT F | 0.31 | 8.9 | ( 53) | TCT S | 0.56 | 24.5 | ( 146) | TAT Y | 0.46 | 10.9 | ( 65) | TGT C | 0.80 | 3.4 | ( 20) |
| TTC F | 0.69 | 20.1 | ( 120) | TCC S | 0.01 | 0.5 | ( 3) | TAC Y | 0.54 | 12.6 | ( 75) | TGC C | 0.20 | 0.8 | ( 5) |
| TTA L | 0.37 | 28.5 | ( 170) | TCA S | 0.19 | 8.4 | ( 50) | TAA * | 1.00 | 3.4 | ( 20) | TGA * | 0.00 | 0.0 | ( 0) |
| TTG L(s) | 0.19 | 14.9 | ( 89) | TCG S | 0.05 | 2.2 | ( 13) | TAG * | 0.00 | 0.0 | ( 0) | TGG W | 1.00 | 3.7 | ( 22) |
| CTT L | 0.37 | 28.5 | ( 170) | CCT P | 0.35 | 11.9 | ( 71) | CAT H | 0.16 | 1.7 | ( 10) | CGT R | 0.88 | 46.5 | ( 277) |
| CTC L | 0.02 | 1.3 | ( 8) | CCC P | 0.00 | 0.0 | ( 0) | CAC H | 0.84 | 8.9 | ( 53) | CGC R | 0.11 | 5.7 | ( 34) |
| CTA L | 0.03 | 2.4 | ( 14) | CCA P | 0.55 | 18.6 | ( 111) | CAA Q | 0.82 | 33.4 | ( 199) | CGA R | 0.00 | 0.0 | ( 0) |
| CTG L(s) | 0.02 | 1.2 | ( 7) | CCG P | 0.09 | 3.2 | ( 19) | CAG Q | 0.18 | 7.2 | ( 43) | CGG R | 0.00 | 0.2 | ( 1) |
| ATT I | 0.43 | 27.7 | ( 165) | ACT T | 0.64 | 34.7 | ( 207) | AAT N | 0.20 | 7.4 | ( 44) | AGT S | 0.04 | 1.8 | ( 11) |
| ATC I | 0.57 | 37.2 | ( 222) | ACC T | 0.04 | 2.2 | ( 13) | AAC N | 0.80 | 30.4 | ( 181) | AGC S | 0.15 | 6.4 | ( 38) |
| ATA I | 0.00 | 0.0 | ( 0) | ACA T | 0.28 | 15.3 | ( 91) | AAA K | 0.84 | 61.4 | ( 366) | AGA R | 0.01 | 0.3 | ( 2) |
| ATG M(s) | 1.00 | 24.0 | ( 143) | ACG T | 0.03 | 1.7 | ( 10) | AAG K | 0.16 | 11.9 | ( 71) | AGG R | 0.00 | 0.0 | ( 0) |
| GTT V | 0.55 | 47.5 | ( 283) | GCT A | 0.54 | 63.9 | ( 381) | GAT D | 0.44 | 24.8 | ( 148) | GGT G | 0.80 | 69.1 | ( 412) |
| GTC V | 0.03 | 2.9 | ( 17) | GCC A | 0.02 | 1.8 | ( 11) | GAC D | 0.56 | 31.4 | ( 187) | GGC G | 0.18 | 15.4 | ( 92) |
| GTA V | 0.35 | 30.2 | ( 180) | GCA A | 0.28 | 33.0 | ( 197) | GAA E | 0.84 | 65.7 | ( 392) | GGA G | 0.02 | 1.3 | ( 8) |
| GTG V | 0.07 | 5.7 | ( 34) | GCG A | 0.16 | 18.8 | ( 112) | GAG E | 0.16 | 12.7 | ( 76) | GGG G | 0.00 | 0.3 | ( 2) |

**Figure 2.**
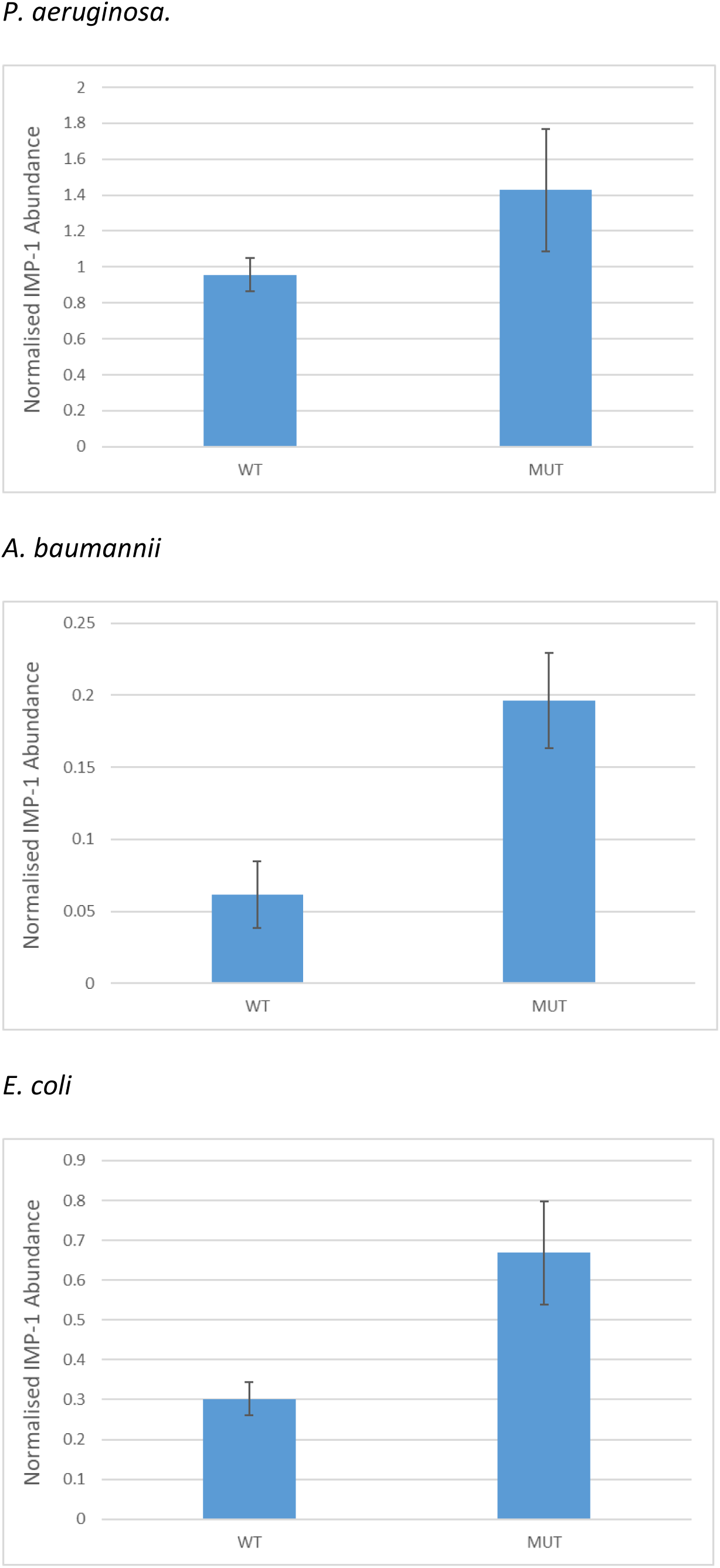
Impact of AAA to AAG lysine codon conversion on IMP-1 protein production. IMP-1 protein abundance in clinical isolates of *P. aeruginosa*, *A. baumannii* and *E. coli* carrying the cloned wild-type *bla*_IMP-1_ gene (WT) and a variant having 17 AAA to AAG mutations (MUT). Protein abundance was measured in sonicated cell extracts and normalised using the abundance of the dominant selectable marker protein for the cloning vector carrying *bla*_IMP-1_. This was, for *E. coli*, where pSU18 was the cloning vector, Cat (chloramphenicol acetyl transferase) and for *P. aeruginosa* and *A. baumannii*, where pUBYT was the cloning vector, AphA (aminoglycoside [Kanamycin] phosphotransferase). Data are means +/-Standard Error of the Mean, *n*=3.

These data show that the simple idea of optimization based on average SCUB – even when that average is taken from highly expressed genes confirmed by proteomics – is rather naïve. The codons in an mRNA affect its folding, which affects its stability and the rate of translation initiation and elongation; local and global charged tRNA levels affect translation elongation rate, and this must be optimised and even varied during the translation of an mRNA to allow accurate protein folding (11-14). Our finding of increased IMP-1 production when AAG lysine codons dominate is not due to relative tRNA abundance since there is only one lysine-tRNA, which recognises both AAA and AAG codons (15). Lysine tRNA/codon specific nucleases have been reported in *E. coli* (16), and it is conceivable that AAA/tRNA interactions preferably promote mRNA cleavage, but the effect we report was seen in three very distinct species, and there is no evidence that AAG/anticodon interactions mean less cleavage, even in *E. coli* (16). The most likely explanation for our findings is a report that duplicate AAA lysine codons lead to ribosomal sliding and increased aberrant translation of an mRNA in *E. coli* (17). We analysed the concatenated sequence data for the genes encoding the 20 most abundant proteins in *E. coli* and found that there are eleven AAA,AAA double lysine codons, comprising 7.8% of all AAA codons. In contrast there are eleven AAA,AAG or AAG,AAA and two AAG,AAG double lysine codons and one AAG,AAG,AAG triple lysine codon found amongst these 20 genes, comprising 26.1% of all AAG codons. This suggests that there is selective pressure for the inclusion of AAG codons preferentially where two or more lysines are encoded together. Within the 17 AAA to AAG mutations made in our lysine codon modified *bla*_IMP-1_ gene (**Figure 3**) one is part of an AAA,AAA double lysine codon, with both codons being converted into AAG in the same mutagenesis step (mutations 6 and 7). To test the specific effects of this mutagenesis step, we measured imipenemase activity in cell extracts of *E. coli* MG1655 transformants carrying pSUHIMP variants having an accumulating number of AAA to AAG mutations, from 1 to 17, starting at the 5’ end of the gene. **Figure 4** shows that carrying *bla*_IMP-1_ with 6 or more mutations gives levels of IMP-1 enzyme activity not significantly different from that provided by pSUHIMP-KV, having all 17 mutations (*p*>0.1). Importantly, introduction of mutations 6 and 7, where IMP-1 enzyme activity significantly increases from basal (p<0.03) is the point at which the AAA,AAA double lysine codon is converted to AAG,AAG. Therefore, based on previously published work using in vitro translation experiments (17), we conclude that there is ribosomal slippage at the AAA,AAA run located in the *bla*_IMP-1_ mRNA, reducing the amount of active IMP-1 protein produced. Breaking up this run with AAG codons means more correct translation and so more IMP-1 enzyme activity. Importantly, we see this effect in all three species tested, despite their divergence. There is no evidence for mutation in the AAA,AAA run in any *bla*_IMP-1_ variant sequence in the Genbank nucleotide sequence database, according to blastn, so the increased IMP-1 enzyme production stimulated by this mutation is seemingly not under strong selective pressure *in vivo*. However, there are two IMP variants where the second lysine codon in this run has been mutated in a non-synonymous way. The most common of these is IMP-22 (18).

**Figure 3.**
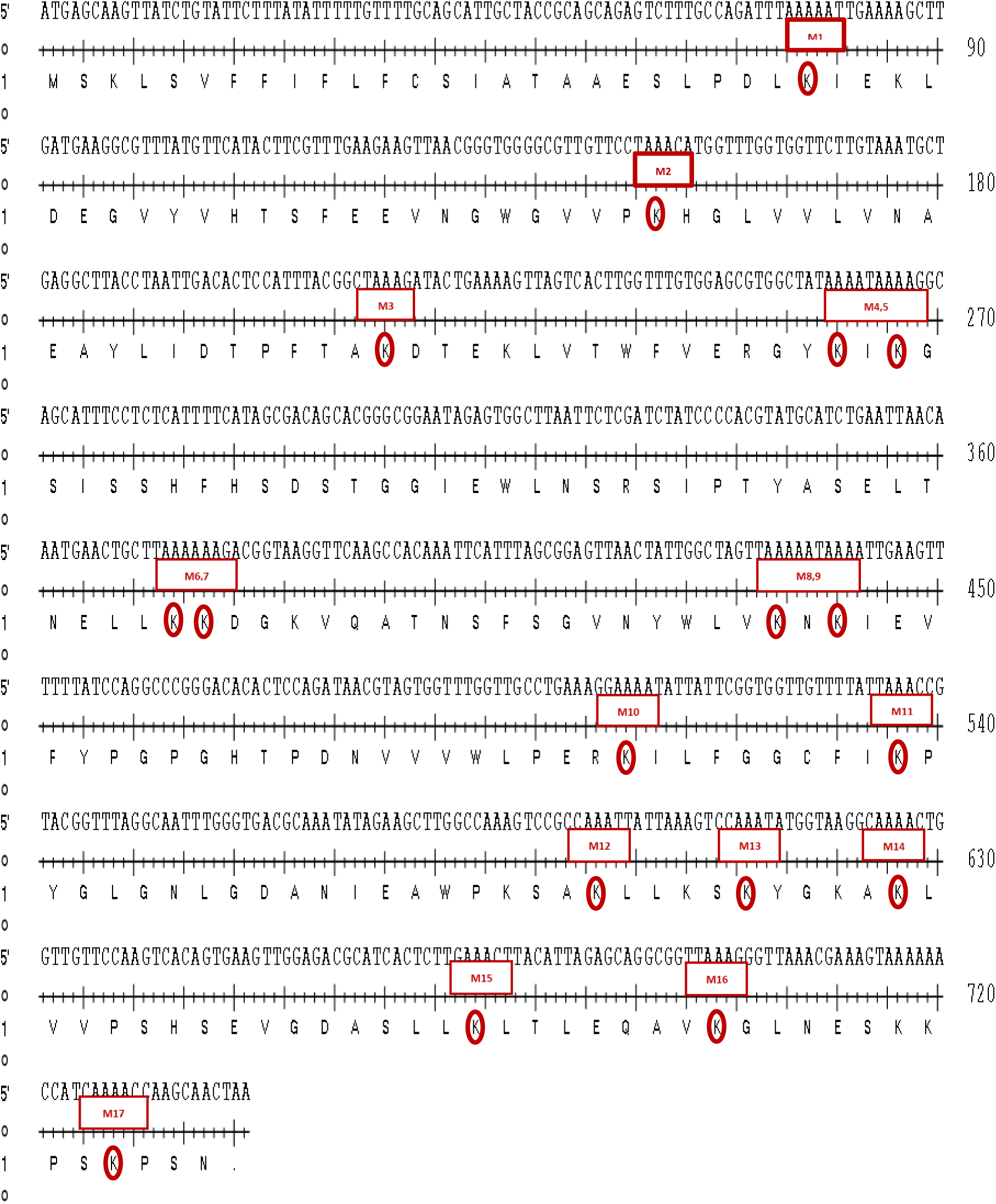
AAA to AAG lysine codon conversions in *bla*_IMP-1_. The 17 AAA lysine codons converted to AAG are marked and sequentially numbered in the blaIMP-1 coding sequence. In some cases, two adjacent mutations were made at using a single primer in the same mutagenesis step, and are labelled as such: M x,y where x and y represent the two sequential mutations.

**Figure 4.**
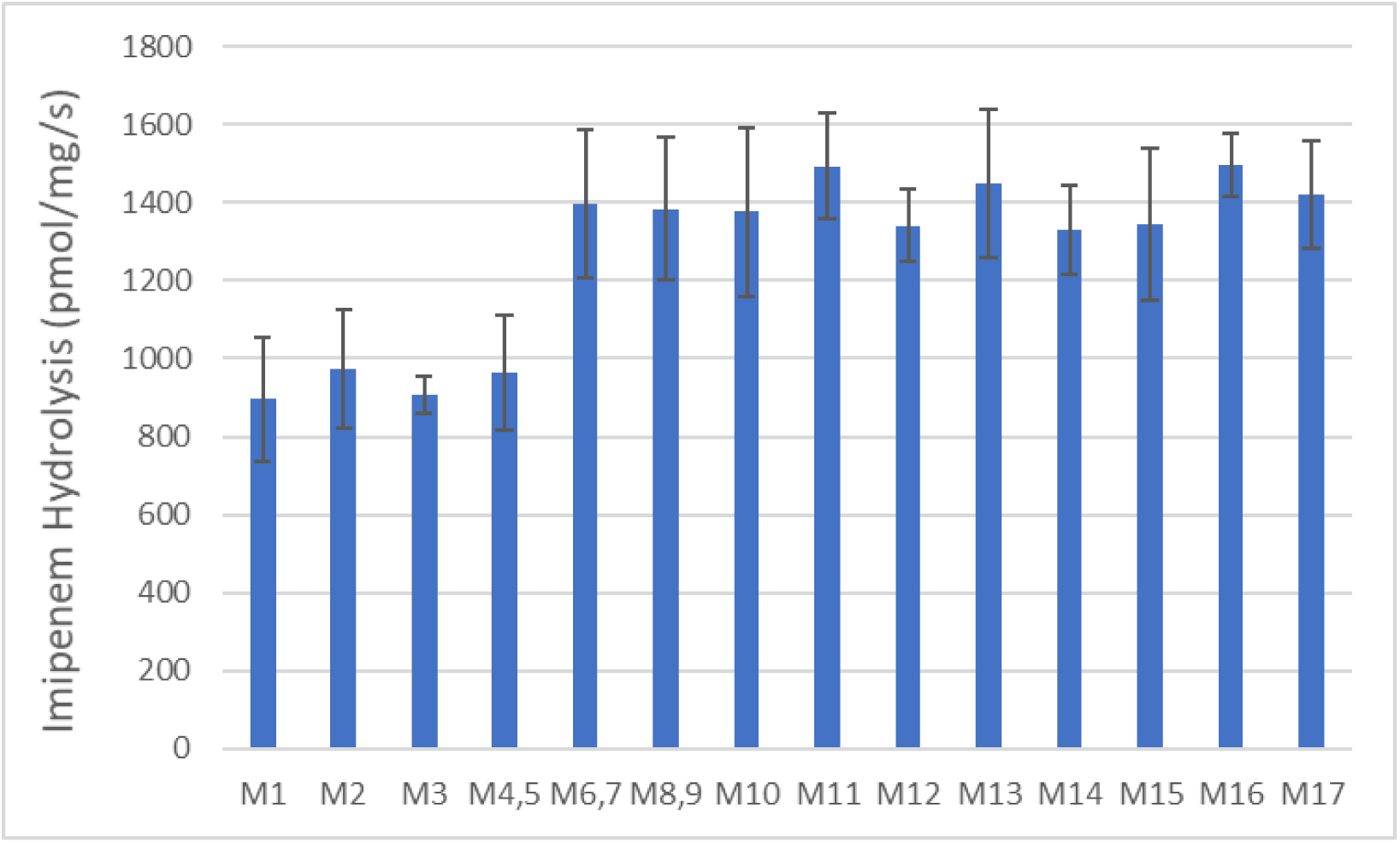
Impact of sequential AAA to AAG lysine codon conversion on IMP-1 production in *E. coli*. IMP-1 enzyme activity was measured using imipenem as substrate in whole cell extracts of *E. coli* MG1655 carrying *bla*_IMP-1_ cloned using pSU18 with one to seventeen AAA to AAG mutations; each mutagenesis step being shown in figure 3. Some steps involved two adjacent mutations, e.g. M4,5. Data are means +/-Standard Deviation, *n*=4.

In conclusion, codon “optimisation” and mutations that change SCUB to be more closely aligned or more distantly aligned to highly expressed genes in multiple bacteria do not guarantee higher levels of gene expression. Indeed, for synonymous lysine codon changes, the increase in IMP β-lactamase production when SCUB was moved further from “optimal” is paradoxical and is most likely to be cause by reduced aberrant protein production that occurs when AAA codons are present in duplicate (17). Care should therefore be taken when interpreting the potential impact of synonymous mutations that affect codon usage in horizontally acquired genes carried on natural plasmids and expressed from native promoters without experimental determination of the effect of these mutations on protein abundance or some phenotypic proxy thereof.

## Experimental

### Bacterial Strains

Bacterial strains used in the study were *E. coli* TOP10 (Invitrogen), MG1655 (19) and a clinical isolate from urine (a gift from Dr Mandy Wooton, Public Health Laboratory for Wales); *P. aeruginosa* PA01 (20) and *A. baumanii* CIP 70-10 (21)

### Molecular Biology

The *bla*_IMP-1_ gene was amplified using PCR. Template DNA was extracted from *P. aeruginosa* clinical isolate 206-3105A (a gift from Dr Mark Toleman, Department of Medical Microbiology, Cardiff University) by suspending a loop-full of bacteria from a fresh Nutrient Agar plate (Oxoid) in 100 μl of molecular biology grade water. The tube was then incubated at 95°C for 15 min and centrifuged at 13,000 rpm for 10 min. The supernatant was removed as a source of template DNA. The integron promoter type upstream of *bla*_IMP-1_ in isolate 206-3105A is PcH1 (22), and there is a *bla*_OXA-2_ gene cassette downstream from bla_IMP-1_ (Genbank accession: AP012280.1). PCR used forward primers which were designed to amplify from the 5’ end of the wild-type PcH1 promoter (5’-ACCCAGTGGACATAAGCCTGTTCGGTTCGTAAACT-3’) into the 5’ end of the *bla*_OXA-2_ gene cassette, (5’-AGCGAAGTTGATATGTATTGTG-3’). Each PCR reaction mixture contained 20 ng of template DNA, 0.4 μmol of each primer, 12.5 μl of RedTaq PCR-ready reaction mix (Sigma-Aldrich) and 8.5 μl of molecular biology grade water. PCR reactions were processed in PTC-100 thermal cycler (Bio-Rad, UK) in 0.2 ml PCR tubes (Starlabs). PCR reaction cycles were 10 min at 95°C, followed by 35 cycles of, 1 min denaturation at 95°C, 1 min annealing at 58°C and 2 min extension at 72°C. The final step was an extension at 72°C for 10 min. The PCR amplicon was TA cloned into the pCR2.1TOPO cloning vector (Invitrogen), removed with EcoRI and ligated into EcoRI linearised RK2-derived vector pRW50 (9) to create the recombinant plasmid pHIMP or the broad host range p15A derived vector pSU18 (23) to create the recombinant plasmid pSUHIMP.

The *E. coli* codon optimized *bla*_IMP-1_ gene variant was designed using the program OPTIMIZER (10) and the variant, including up- and down-stream sequences identical to those seen in pHIMP was synthesized by GeneArt (Thermo-Fisher) and provided, cloned into the cloning vector pMK as the vector pMKH*Ec*IMP. The optimised gene was amplified by PCR using pMKH*Ec*IMP as template and cloned into pRW50, as described for the wild-type gene to create the recombinant plasmid pH*Ec*IMP.

Site directed mutagenesis was performed using the QuikChange^®^ Lightning Site-Directed Mutagenesis Kit (Agilent, UK) according to the manufacturer’s instructions and pSUHIMP as the template. The 17 individual AAA to AAG mutations (**Figure 3**) were introduced in 14 separate mutagenesis steps, each creating a variant with an increasing number of mutations starting at the 5’ end of the gene. The primers were designed using the mutagenesis kit manufacturer’s instructions and are shown in **Table 7**.

**Table 7:**
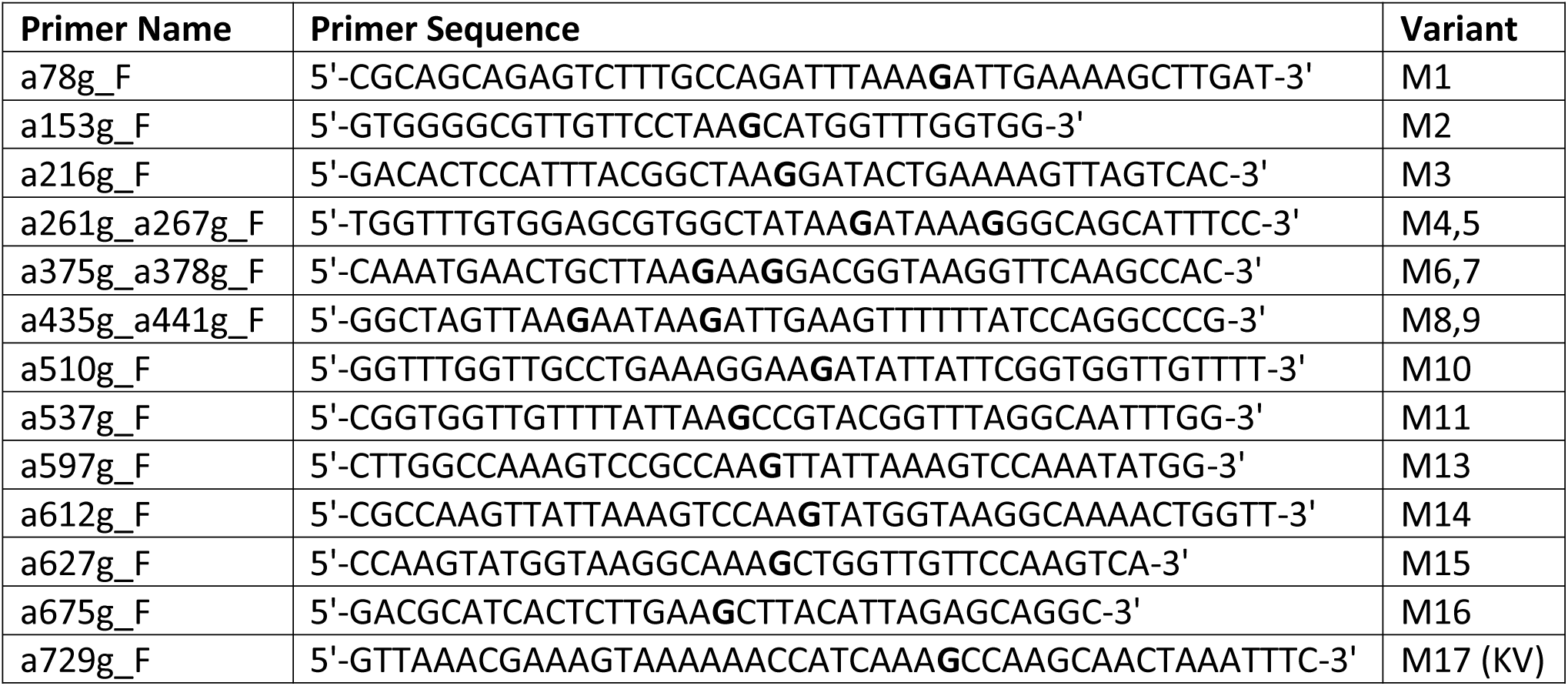
Primers used for Site Directed Mutagenesis

For transformation of *A. baumannii* and *P. aeruginosa bla*_IMP-1_ wild type and variants were subcloned into vector pUBYT being the plasmid pYMAb2 (24) which we modified to remove the OXA promotor region (located upstream of the multiple cloning site) by PCR amplification using primers, 5’-GCAAGAAGGTGATGAATCTACA-3’ and 5’-GTGGCAGCAGCCAACTCA-3’ followed by digestion with XbaI and ligation to produce a circular product.

### Measuring imipenemase specific activity in cell extracts

A volume of 0.5 ml of overnight nutrient bacterial broth culture was added to a 10 ml of fresh nutrient broth which was incubated at 37°C with shaking until an OD_600_ of 0.5-0.6 was reached. The cells were then pelleted by centrifugation at 4,500 rpm for 10 min at 4°C. The pellet was re-suspended in 1 ml of 50 mM HEPES (containing 100 μM ZnCl_2_ at pH 7) and transferred to a tube of lysing matrix B (Fisher Scientific, UK). The cells were lysed using a Ribolyser (Hybaid, UK) at speed of 6.0 for 40 s followed by centrifugation at 13,000 rpm for 1 min to pellet cell debris. The supernatant was used for enzyme activity measurement. Total protein concentration was determined using the Bio-Rad protein assay reagent according to the manufacturer’s instructions. To measure the imipenemase activity in an extract, 100 μl of extract was added to 900 μl of HEPES buffer (containing ZnCl_2_, as above) and 0.1 mM imipenem. Change of absorbance was monitored at 299 nm over 10 min. Specific enzyme activity (pmol imipenem hydrolysed per mg of protein per sec) in each extract was calculated using 7000 M^−1^ as the extinction coefficient of imipenem and dividing enzyme activity with the total amount of protein in each assay.

### mRNA Secondary Structure Prediction

To assess the presence of significant secondary structure in the transcript of wild-type *bla*_IMP-1_ and the *E. coli* SCUB optimized variant, the Mfold program (http://unafold.rna.albany.edu/?q=mfold) was used to predict the folding of the mRNA sequences.

### Preparation of samples from cultured bacteria and proteomics analysis

Bacterial cultures were incubated 50 ml Nutrient Broth (Sigma) with shaking (160 rpm) at 37°C until OD_600_ reached 0.6-0.8. Cells in cultures were pelleted by centrifugation (10 min, 4,000 × g, 4°C) and resuspended in 35 mL of 30 mM Tris-HCl, pH 8 and broken by sonication using a cycle of 1 sec on, 1 sec off for 3 min at amplitude of 63% using a Sonics Vibracell VC-505TM (Sonics and Materials Inc., Newton, Connecticut, USA). The sonicated samples were centrifuged at 8,000 rpm (Sorvall RC5B PLUS using an SS-34 rotor) for 15 min at 4˚C to pellet intact cells and large cell debris and protein concentration in the supernatant was determined using the Bio-Rad Protein Assay Reagent according to the manufacturer’s instructions. One microgram of total protein was separated by SDS-PAGE using 11% acrylamide, 0.5% bis-acrylamide (Bio-Rad) gels and a Bio-Rad Mini-Protein Tetracell chamber model 3000X1. Gels were run at 150 V until the dye front had moved approximately 1 cm into the separating gel. Proteins in gels were stained with Instant Blue (Expedeon) for 5 min and de-stained in water. The 1 cm of gel lane containing each sample was cut out and proteins subjected to in-gel tryptic digestion using a DigestPro automated digestion unit (Intavis Ltd). The resulting peptides were fractionated using an Ultimate 3000 nanoHPLC system in line with an LTQ-Orbitrap Velos mass spectrometer (Thermo Scientific) as previously described (25). The raw data files were processed and quantified using Proteome Discoverer software v1.4 (ThermoScientific) and searched against the UniProt *P. aeruginosa* PA01 database (5563 proteins; UniProt accession UP000002438), the *A. baumannii* ATCC 17978 database (3783 proteins; UniProt accession UP0006737) or the *E. coli* MG1655 database (4307 proteins; UniProt accession UP000000625). The database file is provided as supplementary data. Proteomic searches against the databases was were performed using the SEQUEST (Ver. 28 Rev. 13) algorithm. Protein Area measurements were calculated from peptide peak areas using the “Top 3” method (26) and were then used to calculate the relative abundance of each protein. Proteins with fewer than three peptide hits were excluded from the analysis.

### Codon Usage Calculation

The open reading frames of the 20 most highly expressed genes in each species were downloaded from Genbank and concatenated into a single reading frame. The codon usage calculator (https://www.biologicscorp.com/tools/CodonUsageCalculator#.WrZ02IjFI2w) was applied to this concatenated open reading frame using the standard genetic code table.

## Funding

This work was funded by grant MR/N013646/1 to M.B.A. and K.J.H. from the Antimicrobial Resistance Cross Council Initiative supported by the seven United Kingdom research councils. A

Additional support came from a grant to M.B.A. from the British Society for Antimicrobial Chemotherapy. M.A. and A.M.A were both supported by Postgraduate Scholarships from the Cultural Bureau of the Kingdom of Saudi Arabia.

## Transparency Declaration

None to declare – All authors.

